# Faster 3D saturation-recovery based myocardial T1 mapping using a reduced number of saturation points and denoising

**DOI:** 10.1101/721167

**Authors:** Giovanna Nordio, Aurélien Bustin, Freddy Odille, Torben Schneider, Markus Henningsson, Claudia Prieto, René M. Botnar

## Abstract

**Purpose:** To accelerate the acquisition of free-breathing 3D saturation-recovery-based (SASHA) myocardial T1 mapping by acquiring fewer saturation points in combination with a novel post-processing 3D denoising technique to maintain high accuracy and precision.

**Methods:** 3D SASHA T1 mapping acquires nine T1-weighted images along the saturation recovery curve, resulting in long acquisition times. In this work, we propose to accelerate conventional cardiac T1 mapping by reducing the number of saturation points. High T1 accuracy and precision is maintained by applying a 3D denoising technique to the T1-weighted images prior to pixel-wise T1 fitting. The proposed approach was evaluated on a T1 phantom and 20 healthy subjects, by varying the number of T1-weighted images acquired between three and nine, both prospectively and retrospectively. Three patients with suspected cardiovascular disease were acquired using five T1-weighted images. T1 accuracy and precision was determined for all the acquisitions before and after denoising.

**Results:** In the T1 phantom, no statistical difference was found in terms of accuracy and precision for the different number of T1-weighted images before or after denoising (P=0.99 and P=0.99 for accuracy, P=0.64 and P=0.42 for precision, respectively). In vivo, both prospectively and retrospectively, the precision improved considerably with the number of T1-weighted images employed before denoising (*P*<0.05) but was independent on the number of T1-weighted images after denoising.

**Conclusion:** We demonstrate the feasibility of accelerating 3D SASHA T1 mapping by reducing the number of acquired T1-weighted images in combination with an efficient 3D denoising, without affecting accuracy and precision of T1 values.

## Introduction

Late gadolinium enhancement (LGE) is the reference technique for the visualization of myocardial scar and focal fibrosis, however it cannot visualize diffuse fibrosis (1). In contrast, myocardial T1 mapping allows detection of both focal and diffuse fibrosis and has been extensively investigated as a potential diagnostic tool for the assessment of different cardiomyopathies, such as acute myocarditis, hypertrophic cardiomyopathy, amyloidosis, and dilated cardiomyopathy (2). Several myocardial T1 mapping techniques have been proposed and are based on the acquisition of multiple T1-weighted images along the recovery curve of the magnetization after an initial inversion recovery (3), saturation recovery (4) or a combination of both inversion and saturation recovery (5). The modified Lock-Locker inversion recovery (MOLLI) (3) technique is the most widely used T1 mapping approach, and involves acquiring a single 2D T1 map during a breath-hold (3) or multiple 2D slices during free-breathing (3). MOLLI is highly reproducible and provides precise myocardial T1 maps, however it underestimates T1 due to the acquisition of multiple single-shot T1-weighted images (typically in mid-diastole) spaced over multiple heartbeats after a single inversion pulse (6,7). An alternative technique, the 2D saturation recovery single-shot (2D SASHA) technique, employs a saturation-recovery pulse followed by the acquisition of one T1-weighted image in the same heartbeat, which is then repeated with varying saturation delays (in the order of 9) to sample the entire saturation recovery curve and subsequently generate the myocardial T1 map (4). SASHA permits to obtain more accurate and heart-rate independent T1 maps than MOLLI but its precision is lower, due to the smaller dynamic range compared to inversion recovery based techniques (7,8). Recently, we demonstrated the feasibility of a free-breathing 3D SASHA (9) imaging technique, which allows to provide whole-heart coverage with higher signal-to-noise ratio (SNR) and image resolution than with conventional 2D approaches. The acquisition time of the 3D SASHA sequence is considerably longer than a breath-hold (in the order of 12 min), thus 1D diaphragmatic navigator gating (and tracking) was employed to enable 3D free-breathing T1 mapping (9).

2D and especially 3D T1 mapping techniques suffer from long acquisition times due to the need of acquiring several images along the inversion or saturation recovery curve. Different sampling schemes along the recovery curve have been proposed for 2D MOLLI in order to reduce the number of heartbeats and thus shorten the length of the breath-hold, reducing the total number of heartbeats per breath-hold from 17 to 9 (10). Simultaneous multi-slice imaging techniques have been also investigated, both in free-breathing and under breath-hold, in order to accelerate the acquisition and to increase volume coverage (11,12). However, this approach suffers from limited spatial resolution and does not provide whole-heart coverage. Undersampling reconstruction techniques have also been investigated to accelerate the acquisition and improve the image resolution of T1 maps (13–15).

In this study, we propose to accelerate the 3D SASHA acquisition by reducing the number of the T1-weighted images acquired along the saturation recovery curve. To overcome the expected loss in accuracy and precision due to the reduced number of T1-weighted images acquired, we sought to use a 3D denoising technique based on Beltrami regularization applied directly to the T1-weighted images prior to T1 fitting, enabling accurate and precise 3D SASHA T1 mapping from fewer saturation points and thus shorter scan times. The proposed approach was tested on a standardized T1 phantom, 10 healthy subjects with retrospectively reduced number of T1-weighted images, 10 healthy subjects with prospectively varied number of T1-weighted images and three patients with suspected cardiovascular disease.

## Materials and Methods

### Imaging sequence

The 2D SASHA T1 mapping sequence involves acquiring eleven T1-weighted images at different saturation points (4). The original 3D SASHA T1 mapping sequence previously proposed in (9) acquires and fits nine T1-weighted images along the T1 recovery curve. In this study, the number of images acquired along the saturation recovery curve with the 3D SASHA sequence was varied between three and nine in order to investigate the effect of this parameter on the accuracy and precision of the T1 maps. Fig 1 shows the distribution of three and five time points along the T1 recovery curve used in this study. An image without any saturation preparation was acquired at the beginning of the scan, which corresponds to a measurement of the fully recovered magnetization (last point on the graphs). The saturation time points were then acquired with equal distribution between the minimum and the maximum saturation time (54 ms and 740 ms, respectively, for a heart rate of 60 bpm), following the original implementation of the 2D SASHA (4) imaging sequence.

**Fig 1:**
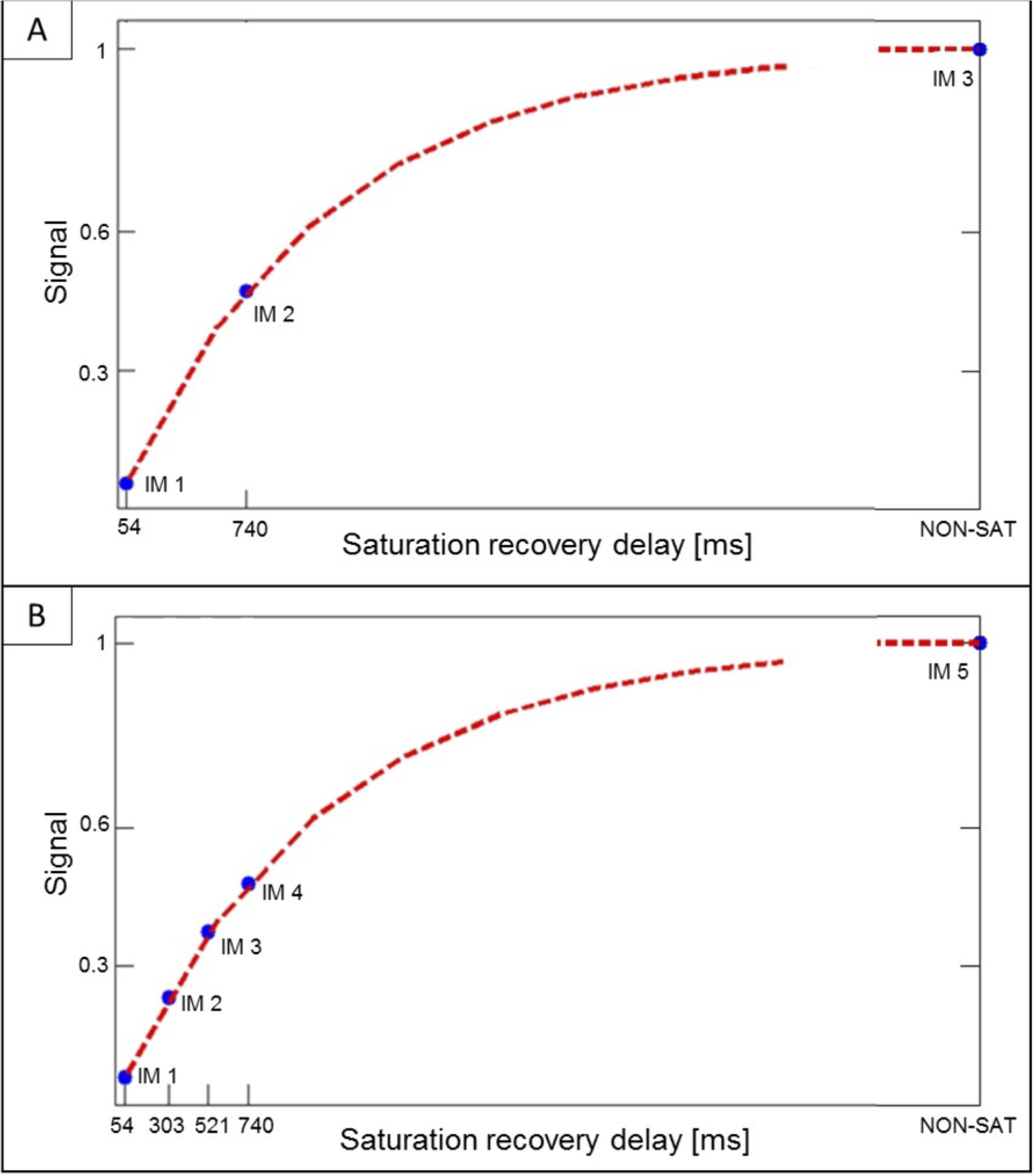
Distribution of three (A) and five (B) time points along the T1 recovery curve.

### 3D Denoising

A reduced number of T1-weighted images is expected to decrease the precision of the corresponding T1 maps, which could result in noisy T1 maps. In order to achieve high precision in spite of the reduced number of saturation recovery time points, we propose to apply a novel 3D Beltrami denoising technique to the T1-weighted images prior to the T1 fitting. The Beltrami framework for image denoising and enhancement was introduced for 2D natural images by Sochen, Kimmel and Malladi (16), proposed for 2D myocardial T1 mapping denoising by Bustin et al (16), and have been recently extended to 3D T1 mapping denoising (17). The Beltrami regularization allows to preserve the edges, while reducing the noise of the images without introducing staircasing artefacts (18).

In this work, the 3D Beltrami denoising framework is applied to 3D T1 mapping with reduced number of time points, by exploiting the redundant information in both the 3D spatial and T1 recovery dimensions, to accelerate the acquisition.

The 3D Beltrami regularization can be expressed as following:

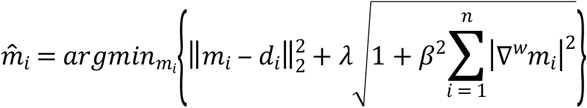

where 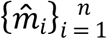 is the 3D set of denoised T1-weighted images for contrast 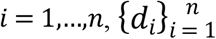 are the corresponding 3D T1-weighted images before denoising, λ is the regularization parameter that controls the trade off between the fidelity to the original acquired data and the Beltrami regularization term, β is the Beltrami constant that allows selection of any arbitrary interpolation between quadratic or total variation gradient penalties and 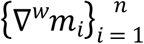 is the 3D weighted-gradient transformation applied to the 3D T1-weighted images with varying contrast. The weighted-gradient function penalizes the gradient depending on its local orientation in order to decrease local smoothness in the directions where sharp transitions are observed, with 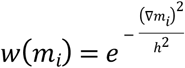 updated at each iteration of the optimization and *h* being the smoothing parameter.

The optimization problem described above was solved using a primal-dual hybrid gradient algorithm (19,20). For all experiments the regularization parameter *λ* was set to 0.25, while the smoothing parameter *h* was set to 5. These parameters were empirically optimized on several dataset (not reported here) to provide the best image quality in terms of SNR and image sharpness. A weight of 1.0 was employed for the Beltrami constant as indicated in (21).

### Imaging

The proposed approach was evaluated in a standardized phantom, 20 healthy subjects and three patients with suspected cardiovascular disease. All imaging studies were performed on a 1.5T MR scanner (Ingenia, Philips, Best, The Netherlands). The study was performed in accordance with the Declaration of Helsinki (2000). All subjects recruited to this study provided written informed consent with study approval from the Institutional Review Board (1/11/12).

### Phantom study

A standardized T1 phantom with nine agar/NiCl2 vials was used for imaging, with T1 values in the range of 250 to 1500 ms (22). The phantom was imaged using the original 3D SASHA sequence, with nine images acquired at different saturation points (54-650 ms). The nominal scan time for the 3D SASHA sequence was of 4:14 minutes:seconds. The number of T1-weighted images for the T1 fitting was reduced retrospectively from three to nine (step size of one). For each case, T1 maps were obtained using a three parameter-fitting model before and after 3D denoising. An inversion recovery spin-echo (IRSE) sequence was used as reference method for the T1 values of the vials. The acquisition parameters used for the 3D SASHA sequence were: FOV = 300 × 300 mm^2^, image resolution = 1.4 × 1.4 mm^2^, slice thickness = 8 mm, flip angle = 35°, TR/TE = 3.3/1.6 ms. Sequence parameters for the spin-echo sequence were: FOV = 200 × 200 mm^2^, image resolution = 3.1 × 3.1 mm^2^, 10 mm slice thickness, TR = 8000 ms and TE = 5.9 ms. The phantom was acquired with a simulated heart rate of 60bpm and using a combination of the 12-channel posterior and 16-channel anterior torso coils.

### Healthy subject study

The effect of reducing the number of T1-weighted images acquired along the recovery curve was studied in healthy subjects retrospectively and prospectively. For the retrospective study, data was collected in 10 healthy subjects using the original 3D SASHA sequence with nine images acquired at different saturation points. The number of T1-weighted images for the T1 fitting was modified retrospectively from three to nine (step size of one) and mapping was performed before and after denoising. For the prospective study, data was acquired in 10 additional healthy subjects using the 3D SASHA sequence with three, five and nine T1-weighted images along the recovery curve. The acquisition parameters used for the 3D SASHA sequence (both retrospective and prospective studies) were: FOV = 300 × 300 mm^2^, image resolution = 1.4 × 1.4 mm^2^, slice thickness = 8 mm, flip angle = 35°, TR/TE = 3.3/1.6 ms, subject specific mid-diastolic trigger delay and saturation times, short-axis orientation, and 32-channel coil. 1D diaphragmatic navigator gating and tracking was used for respiratory motion compensation with a 5mm end-expiratory gating window. For both prospective and retrospective studies, the 3D SASHA T1 map with nine images along the recovery curve was considered as the reference standard. 3D SASHA with nine images along the recovery curve has been previously compared against 2D MOLLI showing excellent agreement in terms of accuracy (17).

### Patient study

The feasibility of using a reduced number of T1-weighted images was investigated in patients using five saturation points along the recovery curve. The number of saturation points for the prospective patient scan was set conservatively based on the results obtained in healthy subjects. 3D SASHA was acquired before contrast injection in three patients with suspected cardiovascular disease with the same parameters used in the healthy subject study but with a lower in plane resolution of 1.6 × 1.6 mm^2^ to further reduce scan time. Late gadolinium enhancement images were acquired 10-20 minutes after injection of 0.1-0.2 mmol/kg of Gadovist and used as ground truth to determine if fibrosis was present. The acquisition parameters used for the 2D LGE sequence include: FOV = 350 × 350 mm^2^, in plane resolution of 1.6 × 1.9 mm^2^, slice thickness = 10 mm, flip angle = 25°, TR/TE = 6.1/3 ms.

### Image analysis

3D denoising and three-parameter fitting were performed offline using MATLAB (MathWorks, Natick, MA). A myocardial T1 map was obtained for each acquisition, using different number of T1-weighted images, both before and after applying the 3D denoising method. A region of interest (ROI) was manually drawn in the septum of the myocardium for the mid slice T1 map in all subjects. Mean and standard deviation in the selected ROI were calculated and used as a measurement of accuracy and precision of the T1 map. The measurements were compared using a Kruskal-Wallis test to identify if there was any significant difference between acquisitions with different numbers of T1-weighted images and between corresponding non-denoised and denoised T1 maps. For statistical analysis, GraphPad Prism v5 for Windows (GraphPad Software, La Jolla, CA) was used with a threshold of *P* < 0.05 to define the statistical significance. A Bland-Altman plot was calculated in order to compare the accuracy and the precision of the 3D SASHA non-denoised vs. 3D SASHA denoised, both for the retrospective and prospective study. In addition, for the prospective study, an AHA segmentation (23) was calculated to compare the accuracy and precision in the different myocardial segments of the whole-heart 3D SASHA for different number of saturation points before and after denoising. This segmentation enables the analysis of the spatial homogeneity of the accuracy and precision for a different number of saturation time points before and after denoising. Three slices (apex, mid and base) and 16 segments were used to represent the cardiac volume, while the 17^th^ segment represented the blood pool.

## Results

### Phantom study

There was good agreement in terms of accuracy between the original 3D SASHA imaging sequence with nine T1-weighted images and the gold standard IRSE sequence, both before and after denoising, as shown in the Bland Altman plot in Fig S1.

Accuracy and precision measured in three phantom vials are shown in Fig 2, with T1 values similar to healthy native myocardium (vial #2), post-contrast myocardium (vial #4) and native blood (vial #6), and for different numbers of T1-weighted images in comparison to the gold standard spin echo values, indicated for each vial in Fig 2. For all the vials, there was no significant difference in terms of accuracy for the different number of T1-weighted images before or after denoising (respectively P = 099 and P = 0.99), as well as in terms of precision before or after denoising (respectively *P* = 0.64 and *P* = 0.42). The nominal scan time for the 3D SASHA sequence with three images acquired along the recovery curve was 1min 25s.

**Fig 2:**
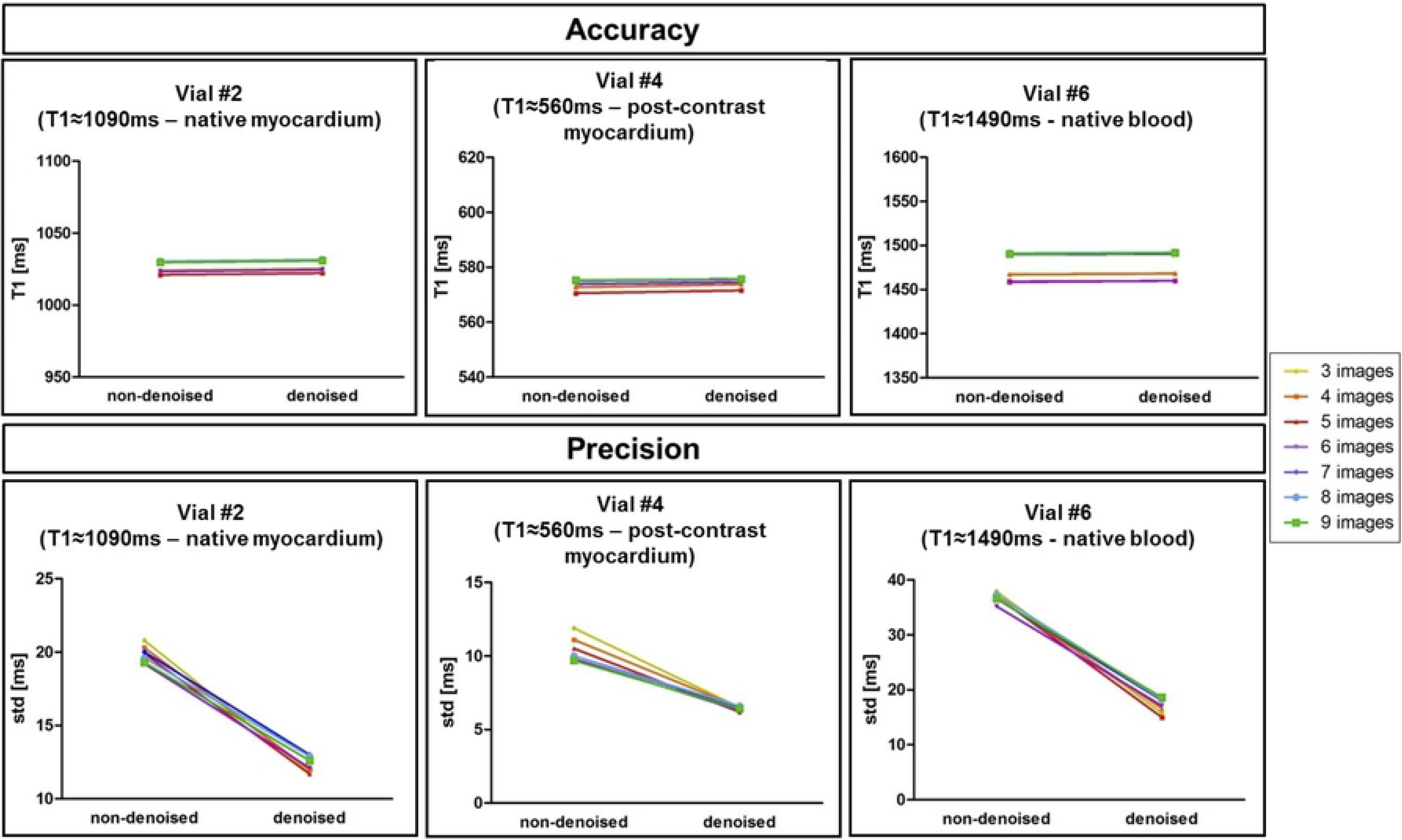
Accuracy and precision of three representative vials of the T1 phantom (native myocardium, post-contrast myocardium, native blood), measured before and after 3D denoising. The number of T1-weighted images used to generate the T1 maps was modified retrospectively from three to nine (step size of one), indicated by the different colors.

### Healthy subjects study

#### Retrospective study

The accuracy and precision averaged over all ten healthy subjects for the retrospective study (three to nine images considered for mapping) both before and after denoising is shown in Fig 3. There was no statistical significant difference between the accuracy measured on the T1 maps reconstructed with three compared to nine T1-weighted images, both before and after denoising (respectively *P* = 0.48 and *P* = 0.14). There was a statistical difference (*P* < 0.05) between the precision measured on the T1 maps reconstructed with three and nine T1-weighted images before denoising, while there was no statistical difference after denoising (*P* = 0.99). There was no statistical difference between the precision measured on the T1 maps reconstructed with four to nine T1-weighted images, both before and after denoising (*P* = 0.99). Fig 4 shows the Bland-Altman plot of the accuracy and precision of the 3D SASHA non-denoised vs. 3D SASHA denoised T1 maps obtained using from three to nine T1-weighted images. The bias indicated in the plot show that there was a slight improvement both in terms of accuracy and precision after 3D denoising when an increased number of T1-weighted images were used.

**Fig 3:**
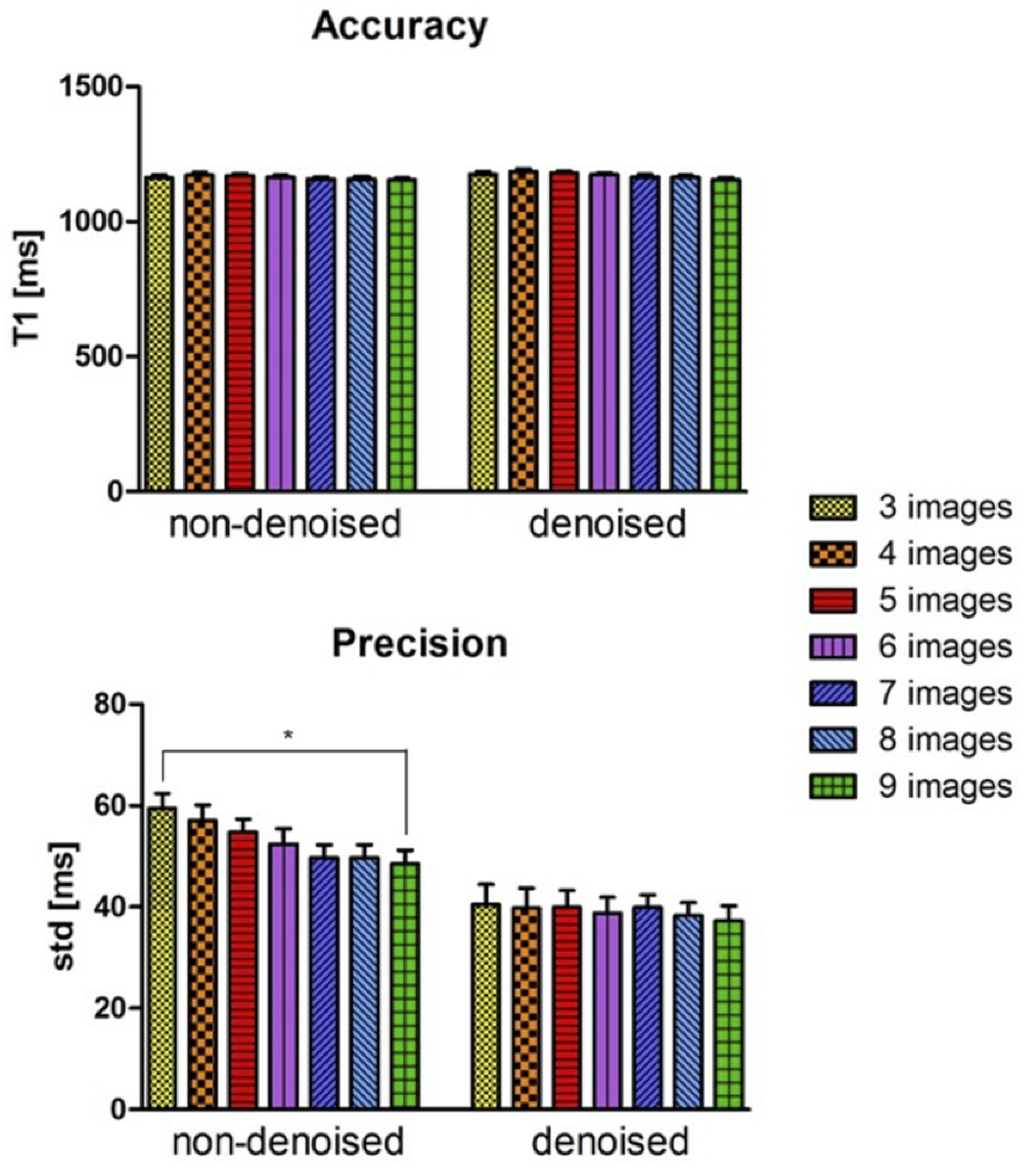
Accuracy and precision averaged between the ten healthy subjects for the retrospective study (3 to 9 images considered for T1 mapping). A ROI was manually drawn in the septum of the myocardium in the mid slice of the 3D SASHA T1 maps before (non-denoised) and after (denoised) denoising. Statistical significance difference is indicated by * (p value < 0.045).

**Fig 4:**
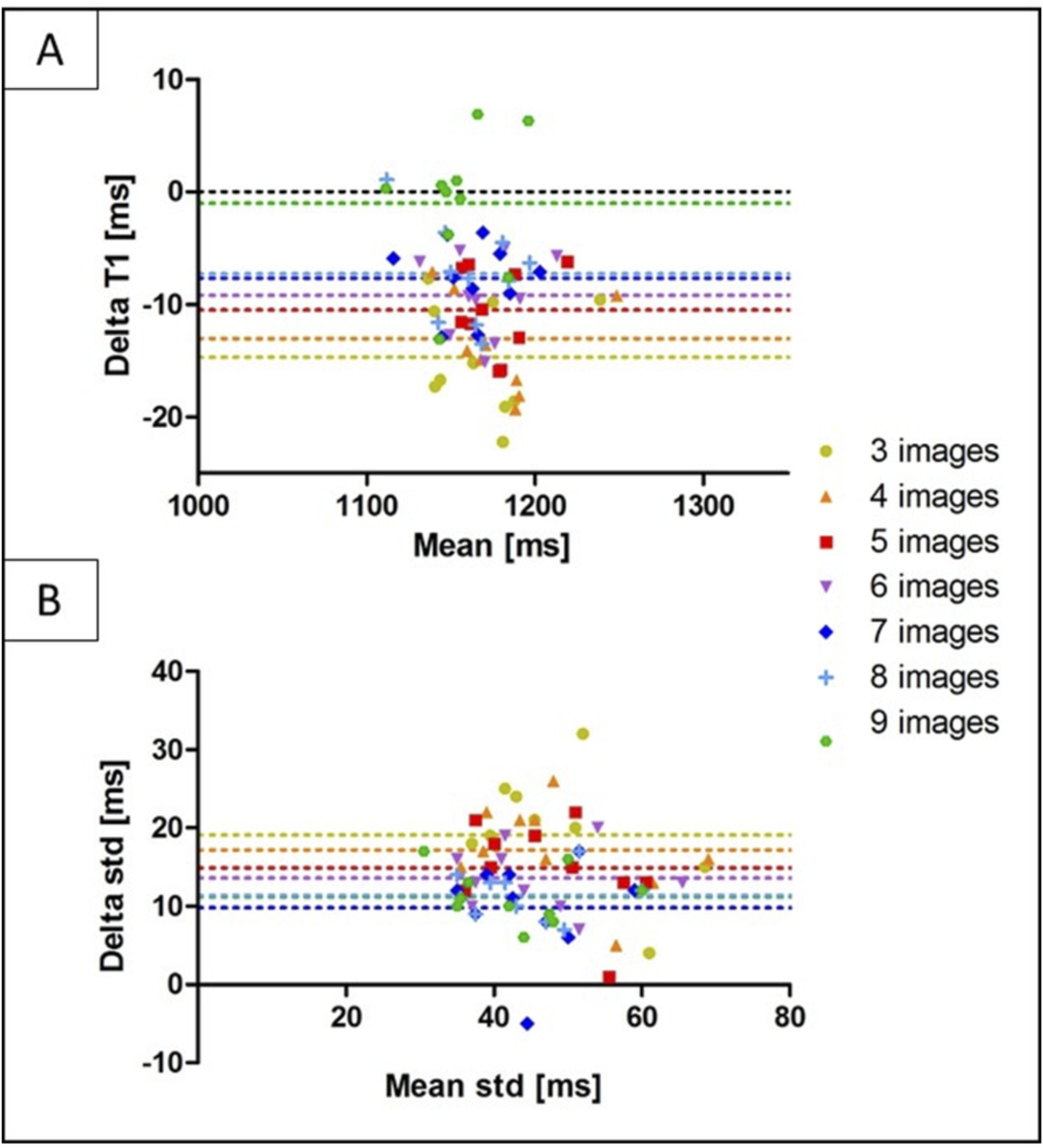
Bland-Altman plot of the accuracy (A) and precision (B) of the non-denoised vs. denoised 3D SASHA using three to nine T1-weighted images for imaging, indicated with different colors in the figure. The bias of each measurement is also indicated in the plot.

3D SASHA T1 maps of two representative healthy subjects, obtained using a different number of saturation points (respectively three, four, five and nine points) before and after denoising are shown in Fig 5. The 3D denoising technique permits to recover the loss in precision due to the lower number of images along the recovery curve and to achieve comparable image quality between the T1 maps reconstructed with a different number of T1-weighted images.

**Fig 5:**
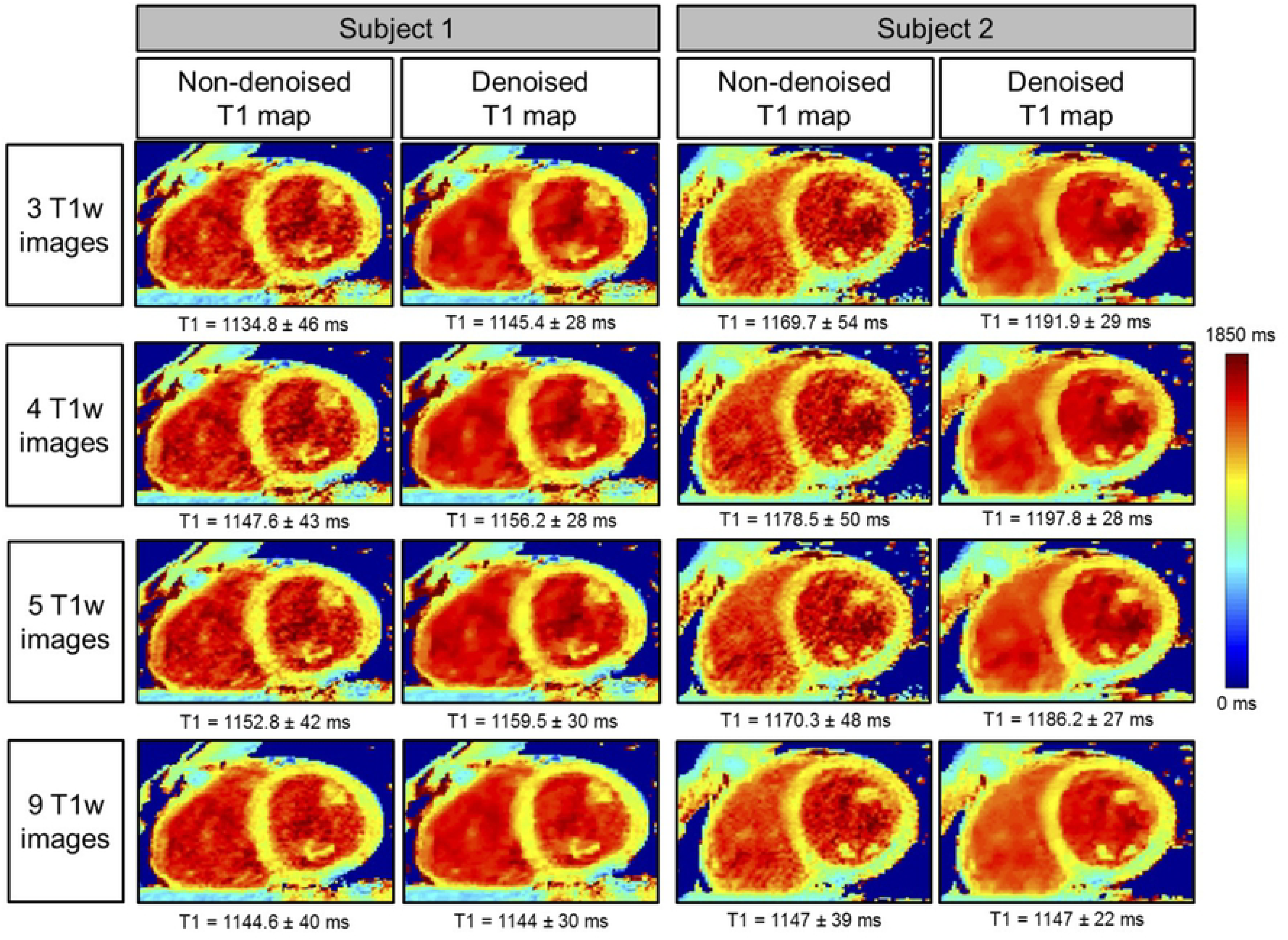
Myocardial 3D SASHA T1 maps of two representative healthy subjects before and after 3D denoising, with the accuracy and precision measured on the corresponding myocardial septum. T1 maps were retrospectively reconstructed by using respectively three, four, five and nine saturation time points along the recovery curve.

#### Prospective study

The accuracy and precision averaged over all ten healthy subjects for the prospective study (three, five and nine T1-weighted images) both before and after denoising is shown in Fig 6a. There was no statistical difference between the accuracy measured on the T1 maps reconstructed using different number of T1-weighted images, both before (*P* = 0.73) and after (*P* = 0.64) denoising. However, a statistical difference (*P* < 0.05) was found between the precision measured on the T1 map acquired with three and nine T1-weighted images before denoising. After applying the 3D denoising technique, the precision was recovered with no statistical difference (*P* = 0.27) when three or nine images where used for T1 mapping. However, if five images were acquired instead of three, the precision improved by about 12%. The average scan time for the original 3D SASHA sequence (nine images along the recovery curve) was 12 ± 1.9 minutes. Scan time was reduced to 7.55 ± 0.7 and 5.6 ± 1.2 minutes when acquiring 5 and 3 images respectively. Fig 6b shows the Bland-Altman plots of the accuracy and precision of the 3D SASHA non-denoised vs. denoised for the three different acquisitions using three, five and nine T1-weighted images. The difference in terms of accuracy between non-denoised and denoised 3D SASHA is smaller when five and nine images are used. In terms of precision, the 3D denoising method has a major impact on the T1 maps acquired with three and five images, with a bias equal to 15.5 compared to 9.6 with nine images.

**Fig 6:**
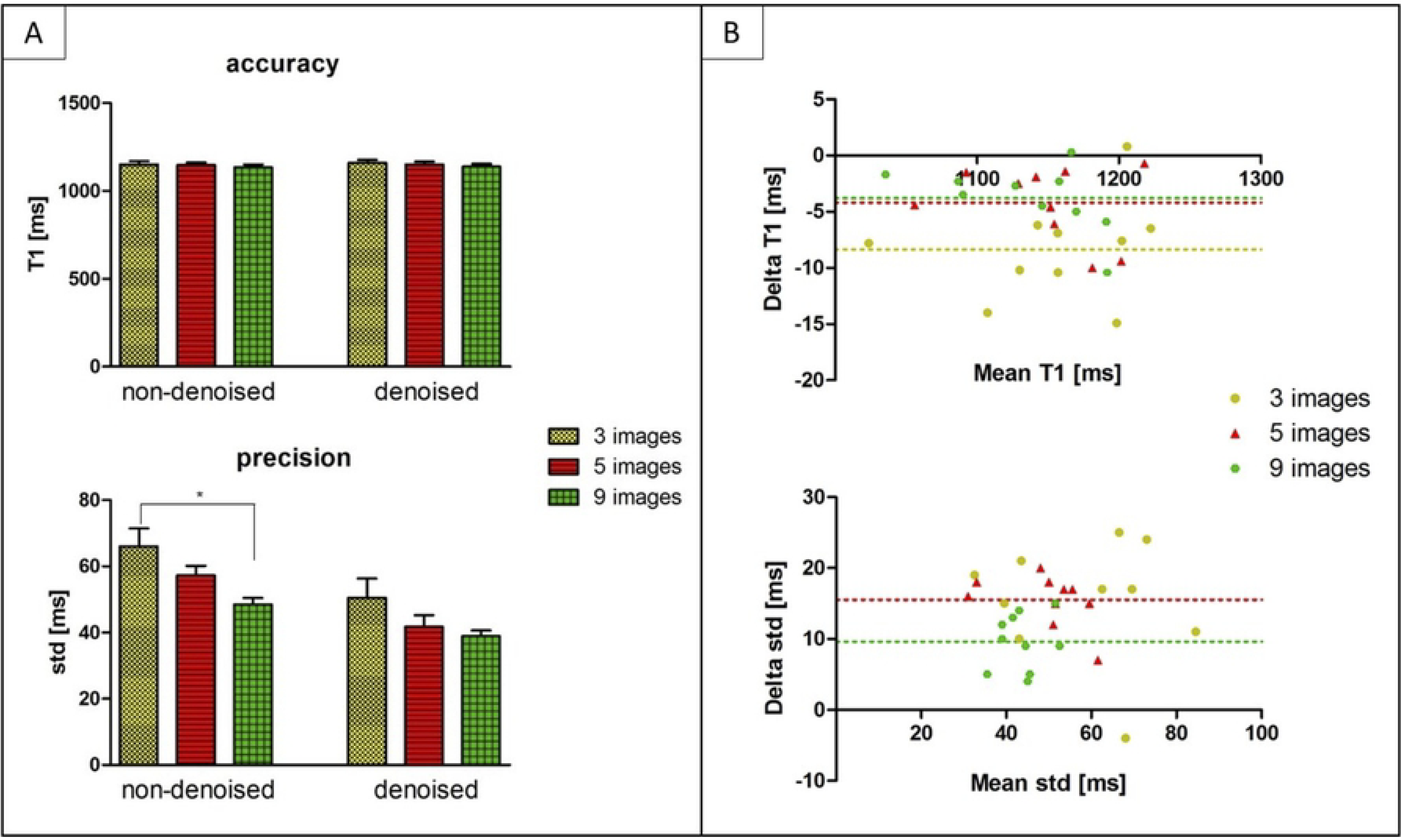
a) Accuracy and precision averaged between the ten healthy subjects, acquired prospectively with 3D SASHA using three (yellow bar), five (red bar) and nine (green bars) saturation time points along the recovery curve. The measurements were performed before (non-denoised) and after (denoised) denoising in the mid slice in the septum of the myocardium. Statistical significant difference is indicated by * (p value < 0.034). b) Bland Altman plot comparing the accuracy (on top) and precision (on bottom) of the 3D SASHA non-denoised vs. 3D SASHA denoised using three, five and nine T1-weighted images. Bias is reported for each graph.

The 3D denoising technique permitted to preserve image quality of the 3D SASHA T1 maps when five T1–weighted images were acquired, as shown in Fig 7 in two representative healthy subjects for the prospective acquisition. However, if only three T1-weighted images were acquired, the delineation of the myocardial borders and the papillary muscles was slightly degraded, as indicated by the white arrows. As the acquisition becomes longer with a higher number of saturation time points, it is also more prone to motion artefacts, as indicated by the blue arrows in the T1 maps for subject 1.

**Fig 7:**
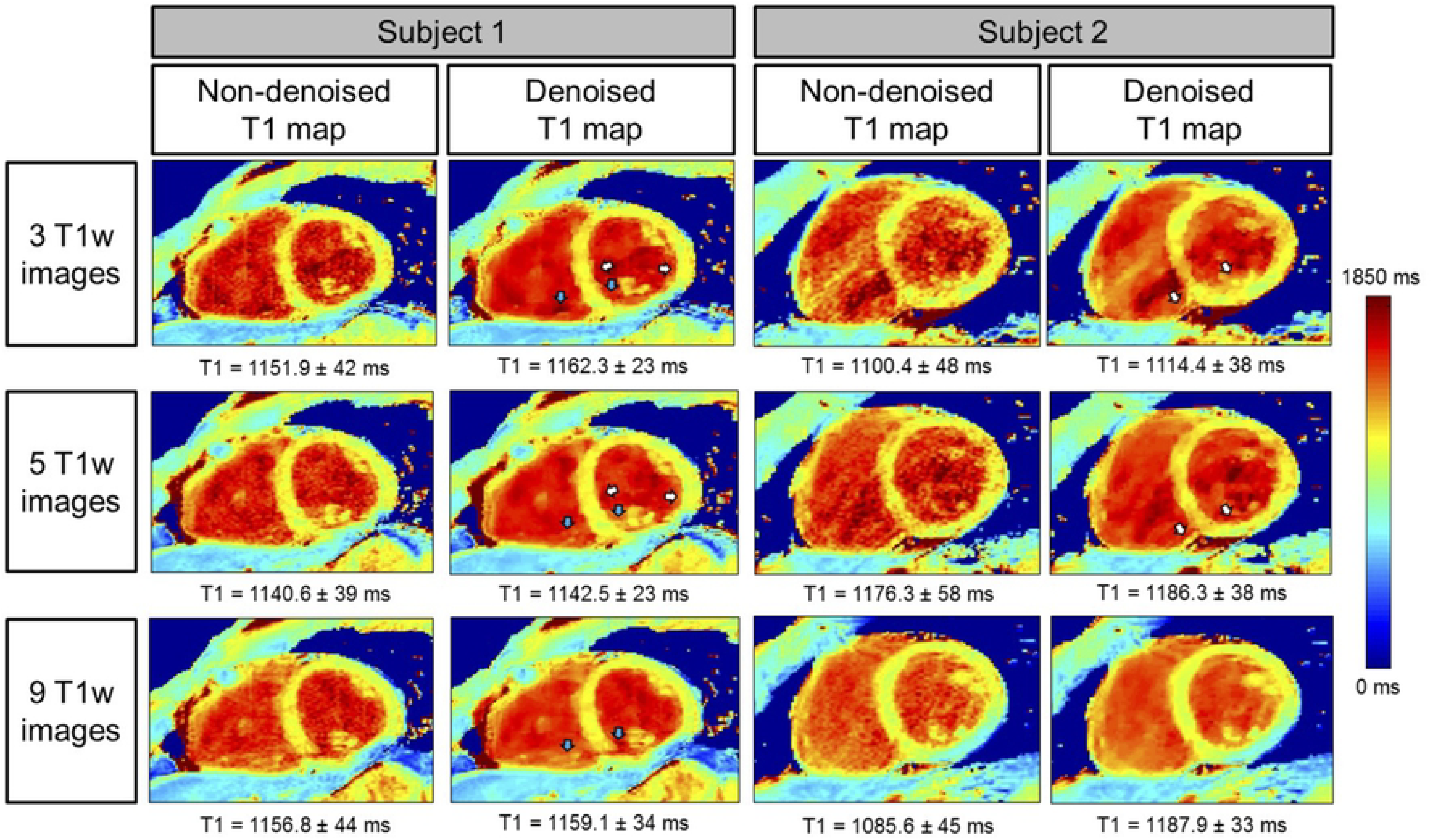
3D SASHA T1 maps of two representative subjects acquired prospectively with three, five and nine T1-weighted images along the saturation recovery curve. The accuracy and precision measured in the myocardial septum are indicated for each T1 map. T1 maps were reconstructed before and after 3D denoising. There was an improvement in the image quality in terms of myocardial and papillary muscles delineation after 3D denoising (white arrows). Motion artefact (blue arrows) can be observed when more images are acquired due to the longer scan time.

Fig 8 shows the AHA segmentation for the accuracy and precision measured on the 3D SASHA T1 maps acquired using three, five and nine T1-weighted images, reconstructed both before and after 3D denoising. The 3D denoising technique does not affect the accuracy of the 3D SASHA T1 maps for all the three different acquisitions (with three, five and nine T1-weighted images). The degree of heterogeneity in percentage are 9.5%, 9.8% and 14% for the denoised 3D SASHA using respectively three, five and nine T1-weighted images. The denoised 3D SASHA T1 maps from three and five T1-weighted images have comparable degree of heterogeneity to the 2D MOLLI presented in Piechnik et al. (10), while with nine T1-weighted images the T1 variability is slightly higher. Instead the precision considerably improves after denoising across the whole left ventricle, particularly when three and five images are acquired.

**Fig 8:**
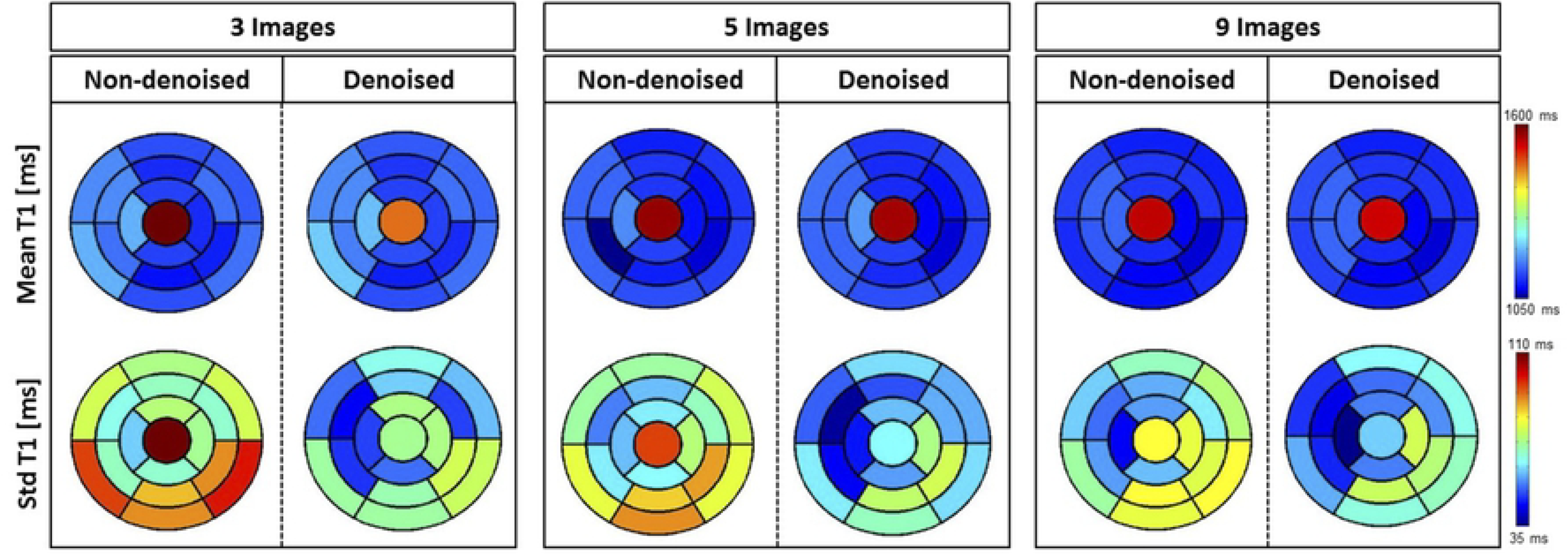
AHA plots for the accuracy (mean) and precision (standard deviation) of the 3D SASHA T1 maps from the ten healthy subjects of the prospective study. The T1 maps were acquired using three, five and nine T1-weighted images and they were obtained both before and after denoising. The accuracy is not affected after denoising, acquiring either 3, 5 or 9 images. Conversely, precision improves after denoising for the three different cases.

### Patient study

Native 3D SASHA T1 maps were obtained in patients using five saturation points prospectively acquired along the recovery curve. T1 maps of the three patients before and after denoising and the corresponding LGE images are shown in Fig 9. Patient 1 was diagnosed with pericarditis, which was detected from both the LGE and the 3D SASHA T1 mapping. The second patient has scar in the apical-mid anterior septum wall. There was an elevation of the T1 values corresponding to the myocardial scar tissue on the 3D SASHA T1 maps, which was in agreement with previously reported myocardial T1 values for scar tissue (24). The third patient, who did not show any cardiac disease, has T1 values in agreement with reference values for native pre-contrast healthy myocardial T1 (25). There was no statistical difference regarding the accuracy and precision measured on non-denoised or denoised 3D SASHA T1 maps (respectively *P* = 0.91 and *P* = 0.19). The average scan time for the 3D SASHA sequence with five images along the recovery curve was about 8 minutes.

**Fig 9:**
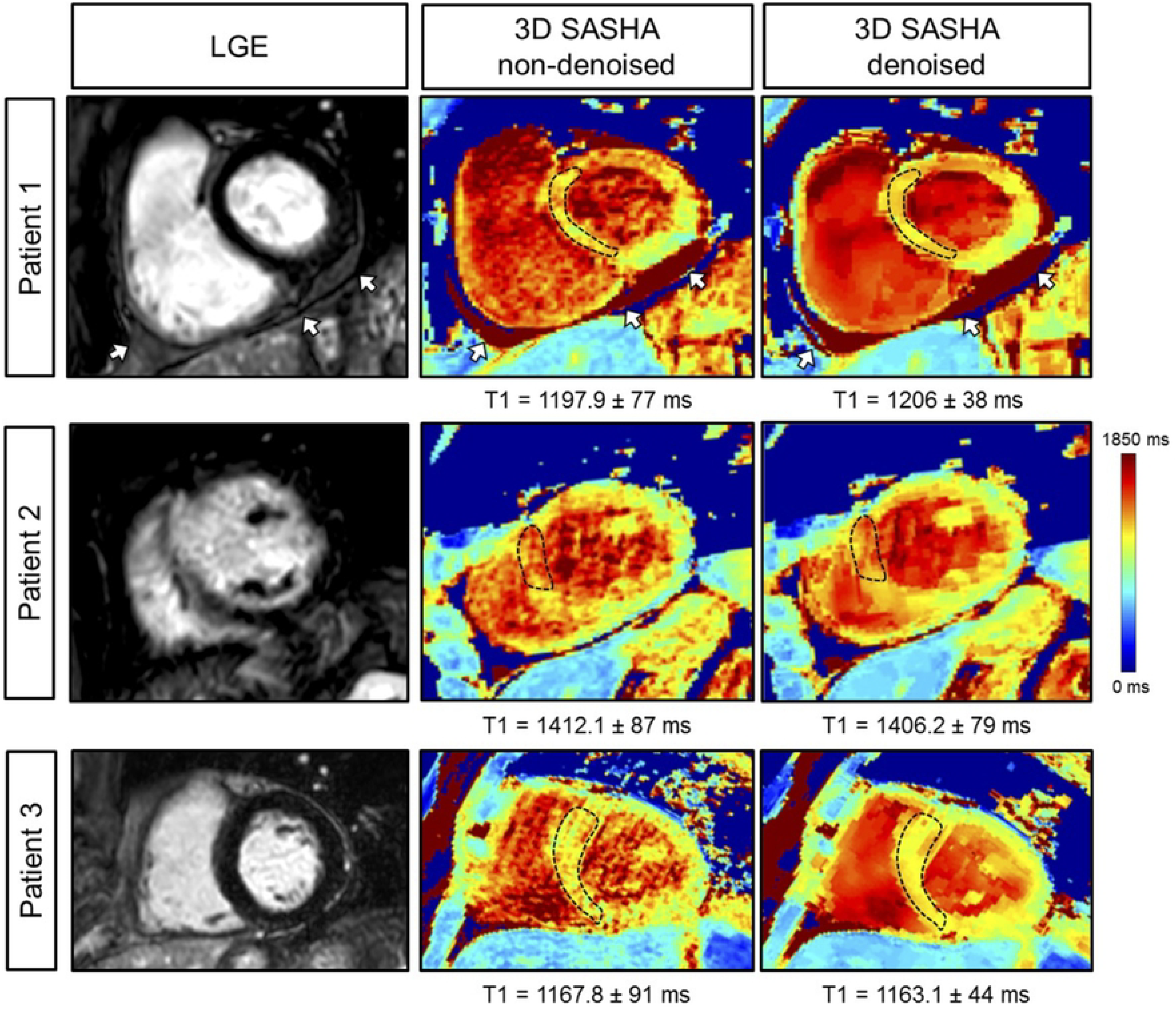
3D SASHA T1 maps before (non-denoised) and after (denoised) denoising and corresponding LGE image for three patients. Patient 1 presented with pericarditis, indicated by the white arrow. Patient 2 had ischemia in the mid-apical septum-anterior wall with corresponding elevated myocardial T1 on the 3D SASHA T1 maps. Patient 3 did not present with disease, which was confirmed by the measured myocardial T1 values.

## Discussion

In this study, we proposed to accelerate the 3D SASHA myocardial T1 mapping technique by reducing the number of saturation time point images acquired along the recovery curve. Our retrospective experiments in a standardized phantom and healthy subjects demonstrate that reducing the number of T1-weighted images used for the T1 fitting has little impact on the accuracy of the T1 maps. However, reducing the number of images has a direct impact on image quality and precision of the T1 maps. To recover the loss in precision and image quality of the myocardial T1 maps when a reduced number of T1-weighted images is acquired, we propose to employ a recently introduced 3D denoising technique based on the Beltrami regularization. The proposed approach was validated prospectively in healthy subjects and feasibility was shown in 3 patients with suspected cardiovascular disease.

The phantom and healthy subjects’ retrospective experiments demonstrated that the accuracy of 3D SASHA T1-mapping was not affected if the number of T1-weighted images considered for T1 mapping was reduced to three images. Moreover, the accuracy was not affected by the application of the 3D denoising technique. Conversely, the precision was reduced when a smaller number of T1-weighted images was employed, with a statistical difference between using three and nine images. No statistical difference was observed in precision after denoising when the number of T1-weighted images was reduced from nine to three. Image quality was affected by the number of time point images used for mapping, which was improved after applying the proposed 3D denoising technique.

For the prospective experiments in healthy subjects, the 3D SASHA was acquired using three, five and nine T1-weighted images along the recovery curve. These experiments confirmed the results obtained in the initial retrospective study: the accuracy of the T1 maps was not affected by the different number of T1-weighted images acquired nor with the application of the 3D denoising technique. There was a statistical difference in precision between the T1 maps acquired with three and nine images before denoising, however no statistical difference was observed after denoising. The AHA segmentation provided a 3D visualization of the accuracy and precision across the whole left ventricle, both before and after 3D denoising. The AHA segmentation confirmed that the accuracy is maintained despite the smaller number of T1-weighted images acquired when combined with the 3D denoising technique. The T1 variability was comparable to 2D MOLLI data previously published in the literature (10) when the 3D SASHA data was acquired with three and five T1-weighted images. The denoised 3D SASHA data acquired with nine T1-weighted images showed higher T1 variability within the left ventricle, probably due to residual respiratory motion. In contrast the precision decreased when a reduced number of acquired T1-weighted images was used for T1 mapping. However, precision improved considerably after denoising. Although not statistically significant, the precision of the T1 map obtained from five T1-weighted images before denoising was about 12% higher than that of the T1 map obtained from three images. In terms of image quality, the delineation of the myocardial borders was in general worst in the T1 maps reconstructed from three images in comparison to the corresponding T1 maps reconstructed from five images (Fig 7). Some respiratory motion related artefacts were observed in the acquisitions with five and nine images consistent with the longer required scan times. In general, a better compromise between image quality and motion related artefacts is observed when the T1 map is reconstructed using five images.

Results from the patient study showed that the proposed accelerated 3D SASHA T1 mapping technique achieves good image quality and provides quantitative values consistent with the complementary LGE images and corresponding diagnosis. The proposed approach enabled a reduction in scan time of about 33% from 12 minutes for the original 3D SASHA technique to approximately ~8 minutes for the proposed accelerated approach with five T1-weighted images along the recovery curve.

The scan time was considerably reduced with the proposed approach by acquiring a smaller number of T1-weighted images. Nevertheless, it was still dependent on the efficiency of the diaphragmatic navigator and its unpredictable scan time, which can severely drop in patients with irregular breathing patterns. Alternative motion compensation techniques, such as image-based navigation and self-navigation (26–28), will be investigated in future studies to achieve 100% respiratory scan efficiency and consequently to further accelerate the scan, which can provide more comfort to the patient and reduce the risk of introducing additional bulk motion artifacts associated with long scans.

The saturation time points were selected along the T1 recovery curve in order to provide an equal distribution between the shortest and the longest saturation time, following the original implementation of the 2D SASHA (4) imaging sequence. Further investigation is warranted to understand the effect of a different selection of the sampling points in terms of accuracy and precision of the T1 map (29).

The 3D Beltrami denoising technique employed in this study permits to considerably improve image quality and the precision of the 3D SASHA T1 maps, independently of the number of T1-weighted images acquired (ranging from nine to three). The denoising is a post-processing step and is applied to the T1-weighted images. Future studies will investigate the integration of the denoising technique as regularization directly in the reconstruction process to further accelerate the scan.

Due to time constraints, only native 3D SASHA T1-mapping was performed in this study. Further studies will investigate the acquisition of accelerated native and post-contrast 3D SASHA myocardial T1 maps in patients with cardiovascular disease to enable the measurement of extracellular volume fraction, which has been demonstrated to be useful to detect diffuse myocardial fibrosis (30). Larger clinical studies are now warranted to validate the clinical value of the proposed imaging technique.

## Conclusions

In conclusion, we have demonstrated the feasibility of accelerating free-breathing 3D SASHA T1 mapping by acquiring fewer (three to five) saturation time point images along the recovery curve. This was achieved by using a 3D denoising method to maintain the high precision of the T1 maps and ensure adequate image quality. The proposed technique permits to acquire a whole-heart high-resolution T1 map in approximately 7 minutes, achieving both high accuracy and precision.

## Acknowledgment

This work was supported by 1) the King’s College London & Imperial College London EPSRC Centre for Doctoral Training in Medical Imaging (EP/L015226/1), 2) EPSRC grants EP/P001009/1 and EP/P007619, 3) the Wellcome EPSRC Centre for Medical Engineering (NS/A000049/1 and WT 203148/Z/16/Z), and 4) the Department of Health via the National Institute for Health Research (NIHR) Cardiovascular Health Technology Cooperative (HTC) and comprehensive Biomedical Research Centre awarded to Guy’s & St Thomas’ NHS Foundation Trust in partnership with King’s College London and King’s College Hospital NHS Foundation Trust.

## SUPPORTING INFORMATION

Fig S1: Bland-Altman plot of the accuracy of the non-denoised (blue) and denoised (red) 3D SASHA vs. the gold standard IRSE sequence, measured in the T1 phantom.

